# A high-throughput screening platform for pigment regulating agents using pluripotent stem cell derived melanocytes

**DOI:** 10.1101/2020.04.10.035295

**Authors:** Valentin Parat, Brigitte Onteniente, Julien Maruotti

## Abstract

In this study, we describe a simple and straight-forward assay using induced pluripotent stem cell derived melanocytes and high-throughput flow cytometry, to screen and identify pigment regulating agents. The assays is based on the correlation between forward light-scatter characteristics and melanin content, with pigmented cells displaying high light absorption/low forward light-scatter, while the opposite is true for lowly pigmented melanocytes, as a result of genetic background or chemical treatments. Orthogonal validation is then performed by regular melanin quantification. Such approach was validated using a set of 80 small molecules, and yielded a confirmed hit. The assay described in this study may prove a useful tool to identify modulators of melanogenesis in human melanocytes.

## Background

The skin pigmentation is largely the result of melanin, a pigment synthesized by melanocytes, specialized cells normally found at the epidermal-dermal junction^1^. Alteration in melanin synthesis may lead to abnormal skin pigmentation, such as hypopigmentation, as is the case with certain forms of albinism^2^. To the contrary, excessive production and accumulation of melanin may cause hyperpigmentation disorders, as observed in melasma, post-inflammatory hyperpigmentation (PIH) or solar lentigines^3^. Hyperpigmentation is a common occurrence in darker skin patients, and treatments often include the use of depigmenting agents. Over the years, a range of high-throughput cell-based assays have been described for screening agents that regulate pigmentation^4–8^. In such assays, human or mouse melanoma cell lines were used for the primary screen. Although immortalized cell lines are amenable to the large scale amplification required for high-throughput screening, they may suffer from abnormal karyotypes^9, 10^ and can display differences with primary melanocytes in terms of behavior^11^ as well as gene^12^ and protein expression patterns^13^. On the other hand, primary melanocytes, especially of human origin, may be difficult to reproducibly obtain in large numbers due to limited cell growth or access to sufficient skin samples.

Human induced pluripotent stem cells (hiPSC) are obtained following the epigenetic reprogramming of donor somatic cells^14^. Since their initial discovery, hiPSC have been differentiated into many cell types^15^, including melanocytes^16^. hiPSC derived melanocytes (hiPSC-MEL) are similar to their primary derived counterparts at the morphological^17, 18^, molecular^17, 19^ and functional levels^16, 19, 20^. Because hiPSC proliferate undefinitely, yet retain their ability to differentiate into any cell lineage, they are suited to large scale production of somatic derivatives, which makes them an appealing alternative to immortalized cell lines for drug screening^21^.

Light scattering analysis by flow cytometry has been showed to correlate with melanin accumulation in melanocytes^22^, as well as with melanosome transfer to keratinocytes in coculture models^23^. Flow cytometers equipped with automated samplers have been developed by several major manufacturers in recent years, and enabled multi-parametric high-throughput analysis in a broad range of applications^24^.

### Question addressed

In this study, we sought to build on the recent advances in both hiPSC and flow cytometry technology. We first confirmed light scatter and melanin content correlation using lowly and highly pigmented hiPSC-MEL. Next, we validated the robustness of this correlation using known depigmenting agents. Finally, we adapted this approach to 96 well plate format and developed a high-throughput flow cytometry (HTFC) screening assay, based on hiPSC-MEL, for the discovery of skin-lightening ingredients.

### Experiment design

#### Cell culture

Human primary melanocytes (HEMn, ThermoFischer Scientific), hiPSC-MEL from african (phototype V-VI) and albinos (OCA1) donors, respectively PCi-MEL_AFR and PCi-MEL_ALB (both from Phenocell) were maintained in a 37°C/5% CO_2_ incubator and cultured on fibronectine coated tissue culture plate (Falcon) in PhenoCULT-MEL (Phenocell). Three days after cell plating, treatment was performed for 4 days with 4, N-Butylresorcinol (BR), arbutin (ARB), hydroquinone (HQ), kojic acid (KA) (all from Sigma-Aldrich), with set 1 of the Stem Cell Differentiation Compound Library or 2-Methoxynaphthoquinone (CAS 2348-82-5) (both from Targetmol).

#### Flow Cytometry

For live cell analysis, cells were dissociated by treatment with TrypLE (Thermofisher Scientific), followed by dilution in Live Cell Imaging Solution (Thermofisher Scientific). For specific marker analysis, cells were stained using the IntraPrep permeabilization kit (Beckman) with 1 μg per million cells of mouse anti-human PMEL17 (Abcam), followed by secondary staining with goat anti-mouse AlexaFluor 488. Data were acquired using an Accuri C6 Plus flow cytometer (BD Biosciences) with auto-sampler.

Additional experimental details are provided in the Supporting Information.

## Results

Both hiPSC-MEL cultures showed the typical dendritic morphology of melanocytes (Fig.1A). Strong pigmentation was observed in PCi-MEL_AFR, while it was nearly absent in PCi-MEL_ALB. Importantly, hiPSC-MEL of both types expressed key melanocyte markers including paired box 3 (PAX3), melanocyte inducing transcription factor (MITF), premelanosome protein (PMEL17) and tyrosinase related protein (TYRP1) in a majority of cells (Fig.1B). Flow cytometry analysis of PMEL17 confirmed its presence in >95% of hiPSC-MEL (Fig.1C). These results are in accordance with previous studies^17, 25^ and show that hiPSC-MEL are pure and express the appropriate specific markers.

**Figure 1.**
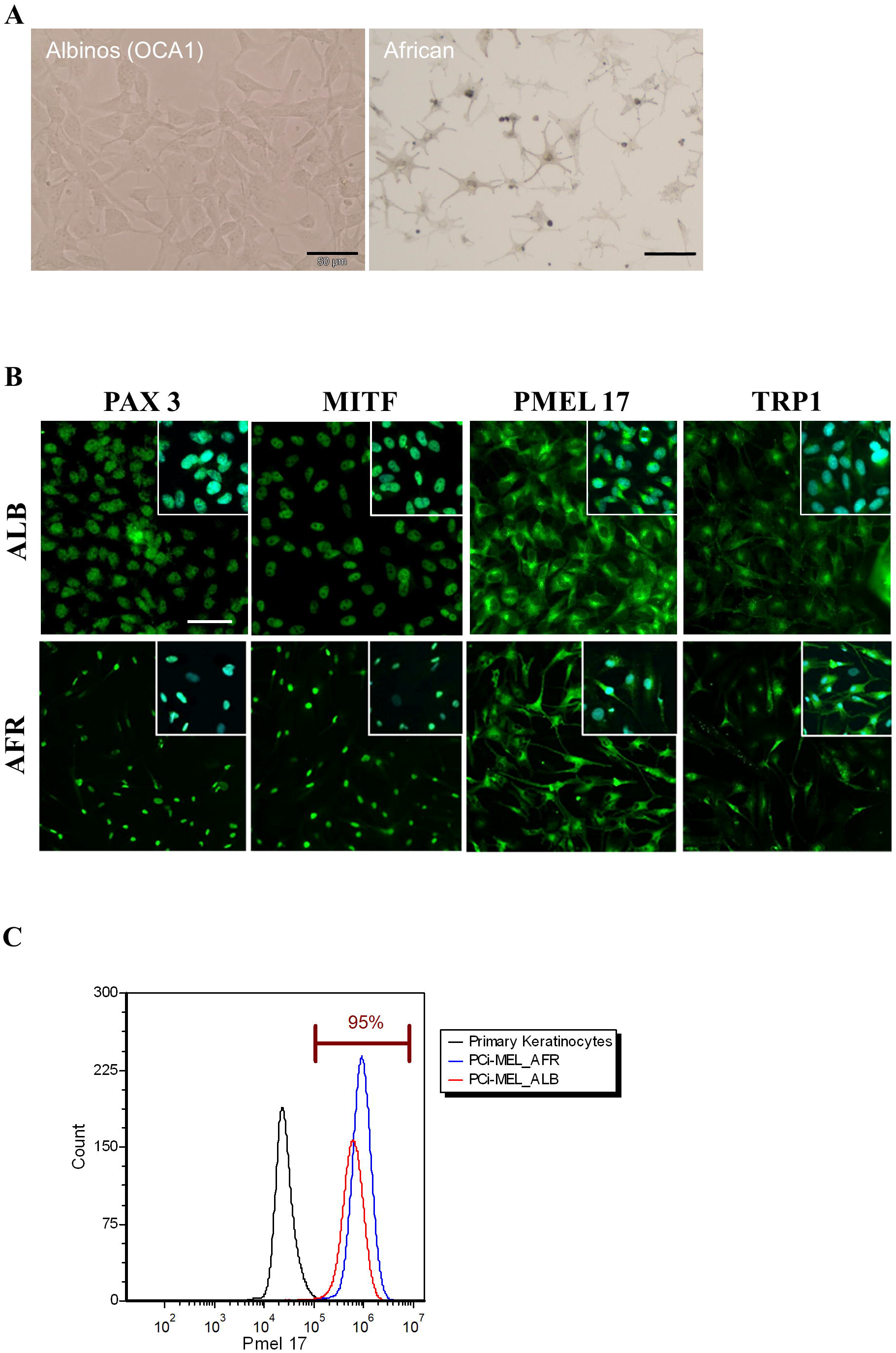
Characterization of hiPSC-MEL. **(A)** Bright field photomicrograph of albinos (OCA1) and darkly pigmented hiPSC-MEL. Scale bars: 50μm. **(B)** Expression of key melanocyte markers (in green) in hiPSC-MEL (Hoescht counterstaining in blue). Scale bars: 100μm. **(C)** Flow cytometric analysis of PMEL17 expression in hiPSC-MEL.

Next, HEMn and hiPSC-MEL were analyzed by flow cytometry for forward light scatter (FSC-A). Previous report^22^ observed that pigmented versus amelanotic melanocytes demonstrated opposite FSC characteristics, as a consequence of high light absorption in melanin rich cells (and conversely low light absorption in melanin deprived cells). Accordingly, pigmented melanocytes, such as HEMn and PCi-MEL_AFR, demonstrated lower FSC-A, while for albinos melanocytes it was much more elevated (Fig.2A). More specifically, median FSC-A for PCi-MEL_AFR was significantly inferior to that of PCi-MEL_ALB by 2 folds, while melanin content followed an opposite trend with PCi-MEL_AFR displaying 20 times more melanin than PCi-MEL_ALB (Fig.2B). Interestingly, median FSC-A for moderately pigmented primary melanocytes HEMn was also significantly above that of PCi-MEL_AFR, in line with melanin levels below that of PCi-MEL_AFR. Treatment with a known depigmenting agent, the tyrosinase inhibitor 4, N-Butylresorcinol (BR)^26^, led to a significant decrease in melanin levels for both HEMn and PCi-MEL_AFR (Fig.2C), while concomitantly increasing FSC-A, compared to vehicle control (Fig.2D). BR treatment did not affect PCi-MEL_ALB median FSC-A, confirming that changes in this parameter correlate with melanin content. Overall, these data demonstrate that median FSC-A can be used to assess pigmentation levels in melanocytes of different origins.

**Figure 2.**
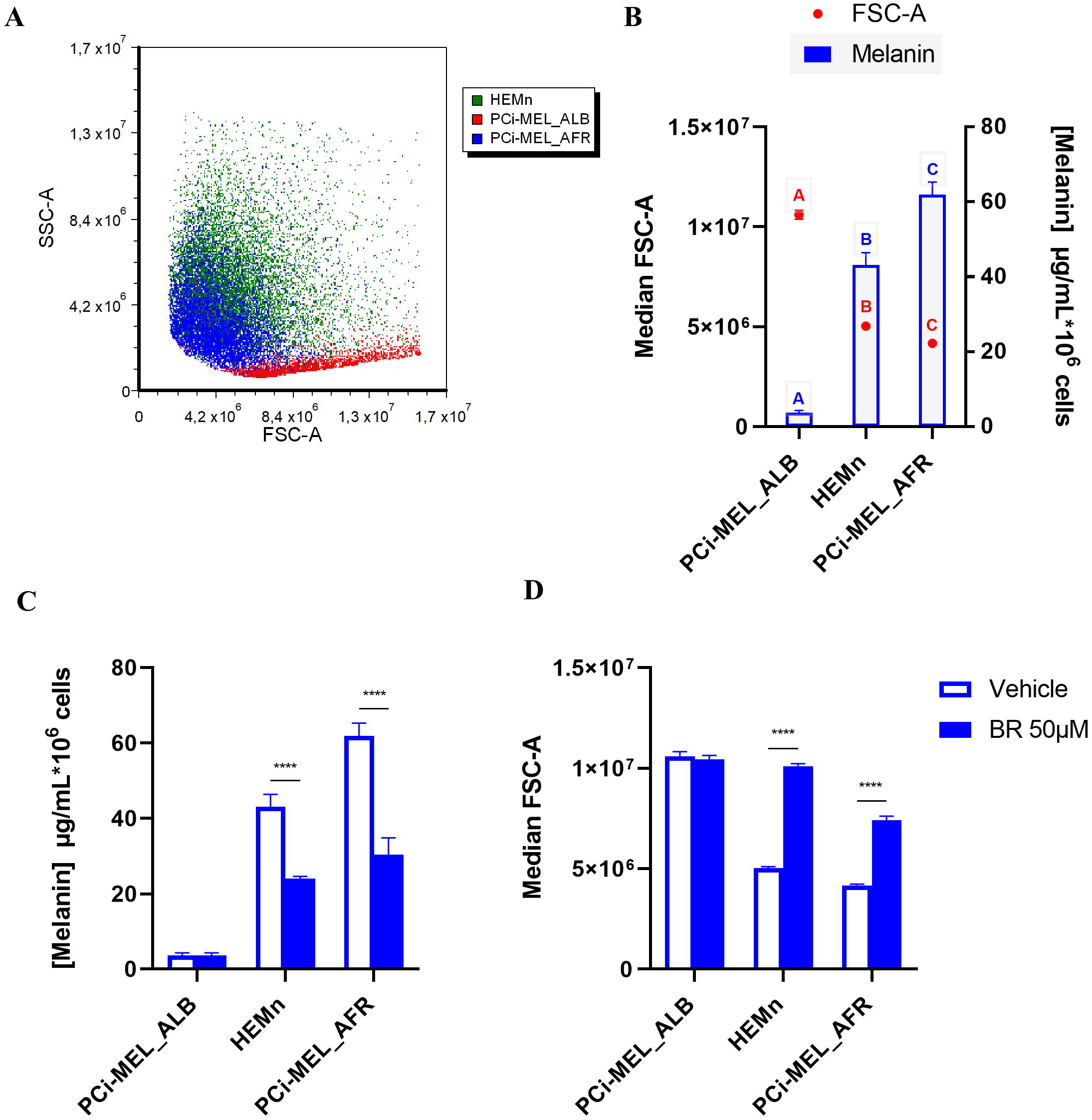
Analysis of forward light scatter and intracellular melanin content in primary and hiPSC derived melanocytes (values represent means ± SD). **(A)** Light Scatter analysis by flow cytometry. **(B)** Intracellular melanin content (blue bars, right axis) and Median FSC-A (red points, left axis). Samples with differing letters are significantly different (by ANOVA). **(C)** Intracellular melanin content and **(D)** Median FSC-A after treatment with melanogenesis inhibitor BR (Butylresorcinol). Vehicle control samples were the same as in (B), (****p<0,0001, by multiple T-test).

We therefore decided to adapt the assay to 96 well plate format, whereby PCi-MEL_AFR were cultured for 3 days, before treatment with compounds for 4 days, followed by HTFC analysis of FSC-A and cell number (Fig.3A). A range of known melanin synthesis regulators were first assessed to validate the assay: as observed in the previous step, BR significantly increased median FSC-A compared to vehicle, which was also true for arbutin (ARB), hydroquinone (HQ) and Kojic Acid (KA) (Fig.3B), with p-value inferior to 0.0001 (by ANOVA, N=8). Whereas cell counting remained unchanged for BR treated samples compared to vehicle, significant cell losses were observed for ARB, HQ and even more so for KA treated wells (Fig.3C). Such effects have already been documented elsewhere^27–29^, and underscore a key aspect of this assay, which can in a single reading assess compound effect on pigmentation and cell survival. On a set of 3 independent plates, the average Z-factor was 0.665, which is within acceptance criteria for high-throughput screening assays (>0.5 being indicative of a good assay)^30^. In order to validate the relevance of this HTFC assay for new pigmentation regulator agent discovery, PCi-MEL_AFR were treated with a set of 80 compounds, each at 10μM in a single replicate. At the end of the treatment, differences in pigmentation were noticeable to the naked eye (Supp. Fig. 1A), although various extents of cell death were observed in some of the depigmented wells (data not shown). Median FSC-A and cell concentration per well were analyzed in parallel (Fig. 3D and 3E). This led to the identification of 2-Methoxynaphthoquinone (MNQ), as the compound with the highest median FSC-A without any detrimental effect on cell survival. PCi-MEL_AFR treated with increasing doses of MNQ displayed a progressive decrease in pigmentation (Supp. Fig.1B). Melanin quantification was used as an orthogonal assay for hit validation (Supp. Fig.1C). It confirmed a dose-dependent response to MNQ, with over 3 fold reduction in melanin synthesis at the 10μM dose (Fig.3F), without adversely affecting cell survival (Fig. 3G). Interestingly, MNQ isolated from Impatiens balsamina has been described previously as a melanogenesis inhibitor^31^, while other naphtoquinone derivatives have also been reported^32^ with such activity^32^. Taken together, these results show that HTFC screening of hiPSC-MEL based on forward light-scatter properties has the ability to identify potent pigmentation regulator agents.

**Figure 3.**
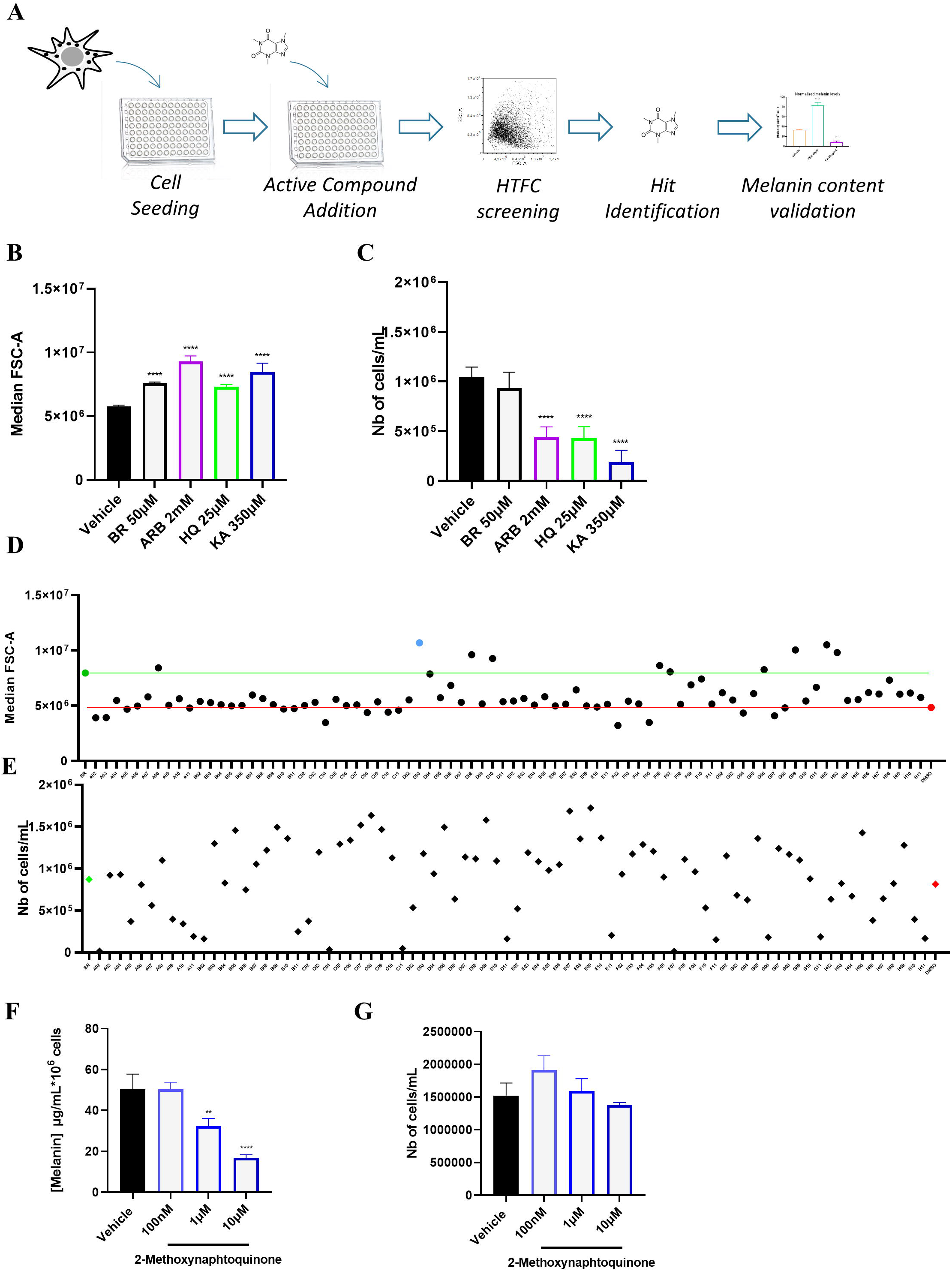
High-throughput flow cytometry assay development and validation (values represent means ± SD). **(A)** Schematic overview of the proposed strategy for HTFC screening of pigmentation regulator agents. **(B)** Median FSC-A and **(C)** cell count after treatment of PCi-MEL_AFR with a panel of melanin modulators such as BR (Butylresorcinol), ARB (Arbutin), HQ (Hydroquinone) and KA (Kojic Acid). (****p<0,0001, ***p<0,001, by ANOVA). **(D)** Median FSC-A and **(E)** cell count after treatment of PCi-MEL_AFR with a panel of 80 compounds (vehicle control in red, BR treated samples in green, and primary hit in blue). **(F)** Intracellular melanin content and **(G)** cell count after treatment of PCi-MEL_AFR with 2-methoxynaphtoquinone (**p<0.01, ****p<0,0001, by ANOVA).

## Conclusion

We describe here a simple yet powerful assay to simultaneously assess pigmentation and cell survival in melanocytes, using HTFC. The assay was validated using known melanin regulators, while a pilote screen of 80 compounds led to the identification of an active ingredient already described as a skin-lightening agent. Because pigmentation is quantified using forward light-scatter, the assay can be easily multiplexed with additional fluorescent probes, allowing multi-parametric analysis.

Although the HTFC data were analyzed to identify melanogenesis inhibitors in this report, it is important to note that the same assay can also lead to the discovery of pro-melanogenic agents: some of the tested compounds actually displayed a median FSC-A lower than that of the vehicle, suggesting increased melanin content in the melanocytes. Our HTFC screen could therefore be performed in the context of skin tanning, using lightly pigmented hiPSC-MEL. Alternatively, this assay could also prove useful to discover drug candidates in the context of skin hypopigmentation disorders: additional pluripotent stem cell lines have been derived over the past years from oculocutaneous albinism affected donors^16, 33, 34^, while small molecule treatment may have the potential to improve skin pigmentation in such patients^35^.

## Supporting information

Supporting Mat&Met and Supporting Fig legends

---

**Supplementary Figure 1.**
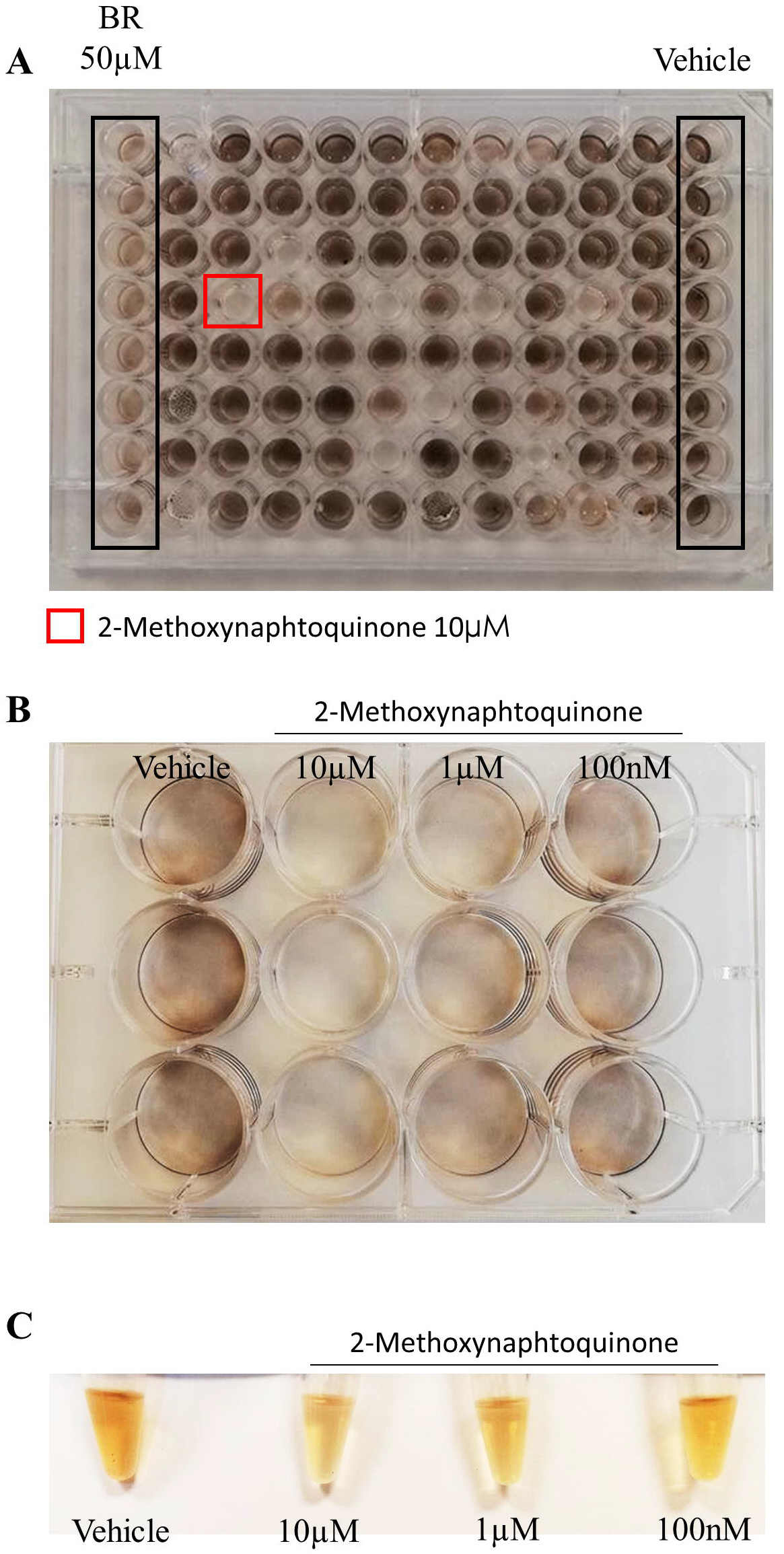

