## Supporting Mat&Met and Supporting Fig legends for "A high-throughput screening platform for pigment regulating agents using pluripotent stem cell derived melanocytes"

**Supporting Information**

**Experiment design**

Immunostaining

Cells were fixed and permeabilized, before staining with mouse anti-human PAX3 (Abcam), mouse anti-human MITF (Thermofisher Scientific), mouse anti-human PMEL17 (Abcam) or mouse anti-human TYRP1 (Abcam). Secondary staining was performed with goat anti-mouse conjugated to AlexaFluor 488 (Thermofisher Scientific), and counterstaining with Hoechst 33342 (Sigma-Aldrich). Microphotographs were taken with an automated microscope (CellInsight, Thermofisher Scientific).

Melanin quantification

Cells were lysed by treatment with NaOH 1M and heated at 80°C for 90 minutes. Absorbance was then measured at 405nm using a Varioskan plus multimode plate reader (ThermoFicher Scientific), and melanin concentration determined against a standard curve from synthetic melanin.

Statistical analysis

Unless otherwise mentioned, experiments were performed on N=3 biological repeats, and data were analysed by multiple T-Test or ANOVA, as appropriate, using Graphpad (Prism) and a 0.05 significance threshold.

**Supplementary figure legends**

**Figure 1. Macroscopic view of PCi-MEL_AFR assay plates and melanin extraction**

**(A)** 96 well plate following treatment with small molecule panel. **(B)** Culture plate following treatment with 2-Methoxynaphtoquinone. **(C)** Cell lysates for melanin quantification by optical density.
